# Mitochondrial Dysfunction Induces Hepatic Lipid Accumulation and Inflammatory Responses in mtDNA Mutator Mice

**DOI:** 10.1101/2025.04.25.650453

**Authors:** Jaime M. Ross, Angela Kim, Kayla Berthod, Rui M. Branca, Maria Olin, Israel Pichardo-Casas, Janne Lehtiö, Ingemar Björkhem, David A. Sinclair, Lars Olson, Giuseppe Coppotelli

## Abstract

**Background:** Non-Alcoholic Fatty Liver Disease (NAFLD), now referred to as Metabolic Dysfunction-Associated Steatotic Liver Disease (MASLD), is a widespread and complex health issue characterized by the accumulation of lipids in liver cells, not linked to alcohol consumption or other known liver problems. Despite its strong association with insulin resistance and obesity, a notable proportion of lean individuals develop NAFLD, suggesting the involvement of additional factors. Mitochondrial dysfunction has emerged as a potential contributor, given its role in lipid homeostasis and the generation of reactive oxidative species (ROS).

**Methods:** In this study, we took advantage of the mtDNA mutator mouse model, characterized by oxidative phosphorylation deficiency due to accelerated accumulation of mitochondrial DNA (mtDNA) mutations, to investigate how mitochondrial dysfunction affects liver homeostasis triggering fat accumulation.

**Results:** We found that mitochondrial dysfunction induces liver steatosis and inflammation in this mouse model at a young age, independent of obesity. Using quantitative mass spectrometry, we reveal that mitochondrial dysfunction alters the levels of multiple proteins involved in lipid metabolism, cholesterol homeostasis, and inflammation. In vitro data show a shift in cellular metabolism toward the glycolytic pathway and confirm the upregulation of inflammatory genes. These changes are associated with oxidative stress and occur independently of mtDNA molecule release into the cytoplasm.

**Conclusion:** Our findings demonstrate that liver steatosis might develop as result of mitochondrial dysfunction without obesity and insulin resistance, underscoring the importance of mitochondrial dysfunction in NAFLD development in lean individuals and providing valuable insights into the molecular mechanisms underlying this complex metabolic disorder. This mouse model offers a unique platform for further investigations aimed at unraveling the intricacies of NAFLD pathogenesis and potential therapeutic targets beyond conventional risk factors.

## Introduction

Non-Alcoholic Fatty Liver Disease (NAFLD), now referred to as Metabolic Dysfunction-Associated Steatotic Liver Disease (MASLD), is a widespread and complex health issue characterized by the accumulation of lipids in liver cells, not linked to alcohol consumption or other known liver problems (1). NAFLD encompasses a range of conditions, from liver steatosis, defined as the buildup of lipid droplets in hepatocytes, considered the earliest stage of NAFLD, to steatohepatitis characterized by inflammation, and, at its most serious, cirrhosis and hepatocarcinoma (2). Approximately 30-40% of adults in the United States are diagnosed with NAFLD, and about 25% of them will progress to nonalcoholic steatohepatitis (NASH), affecting 3-12% of adults (3,4). Symptoms of these diseases often do not appear, but if they do, they may present as fatigue, abdominal pain on the right side, a decrease in appetite with weight loss, swelling of the legs, or easy bleeding/bruising (5). Although NAFLD is strongly associated with insulin resistance and obesity, a significant number of lean individuals develop NAFLD, even in the absence of insulin resistance, type 2 diabetes (T2D), and related metabolic comorbidities (6). This suggests that other factors, such as dietary composition, lifestyle factors, and genetic susceptibility, may contribute to the development of NAFLD. Several reports have indicated that NAFLD might be a mitochondrial disease, although the molecular mechanisms underlying such a causative effect have not been completely understood (7–9).

Mitochondria are key regulators of lipid homeostasis and the main source of reactive oxidative species (ROSs), which are responsible for lipid peroxidation, cytokine release, and cell death (10,11). Several animal models of mitochondrial dysfunction, with and without a high-fat and high-fructose diet, have been used to elucidate the involvement of oxidative phosphorylation (OXPHOS) and β-oxidation impairment in the onset and progression of NAFLD. For example, it has been shown that impairment of β-oxidation due to the removal of the mitochondrial trifunctional protein (MTP), which catalyzes long-chain fatty acid oxidation, induces NAFLD in heterozygous mice (MTP+/-) during normal aging (12,13). In a different mouse model of mitochondrial dysfunction due to the removal of the mitochondrial fission factor (MFF), an NAFLD phenotype was observed after challenging mice with a high-fat diet, but not in mice fed regular chow (14). Mitochondrial DNA mutations, deletions, and haplotype have also been correlated with NAFLD, although the underlying molecular mechanism still remains to be fully elucidated (15–18).

In this study, we investigated the onset and progression of an NAFLD-like phenotype in the mtDNA mutator mouse, a mouse model of OXPHOS deficiency due to the accumulation of mtDNA mutations (19,20). The mtDNA mutator mouse is a homozygous knock-in mouse expressing a proofreading-deficient version of the nucleus-encoded catalytic subunit of mtDNA polymerase-γ (PolgA), which contains an alanine instead of the critical aspartate residue of the second exonuclease domain (D257A) (19). The mtDNA mutator mouse develops high levels of point mutations and linear deletions and shows multiple signs of human-like aging occurring prematurely, including reduced lifespan (42-44 weeks), weight loss, alopecia, anemia, kyphosis, osteoporosis, sarcopenia, and loss of subcutaneous fat. Here, we report the onset of liver steatosis in the mtDNA mutator mouse starting at 25 weeks of age associated with dysregulation of cholesterol metabolism and inflammation. Notably, the NAFLD-like phenotype in this mouse model is not associated with obesity or insulin resistance and could be used to better understand the molecular mechanisms underpinning NAFLD development in lean people (21).

## Materials and Methods

### Animals

Homozygous mtDNA mutator (PolgA^mut/mut^; B6-Polg*^tm1.1(D257A)Lrsn^*) mice, herein referred to as Mut mice, and “wild-type” (PolgA^wt/wt^) nDNA^wt^/mtDNA^mut^ littermates that have a wild-type nuclear genome but maternally inherited mtDNA mutations (22,23), herein referred to as WT mice, were obtained by crossing mice heterozygous (PolgA^wt/mut^) for the mtDNA mutator allele. Animals of both sexes were used for the study and were genotyped as previously described (19). Both mtDNA mutator and WT mice received a standard diet (R34, Lactamin/Lantmännen, Stockholm, Sweden). Three- and 24-month-old C57BL/6J mice were obtained from the NIA Aged Rodent Colonies (Charles River Laboratories, Kingston, NY or Raleigh, NC, USA), were acclimatized at least for a month prior to experimentation, and received a standard diet (5053 - PicoLab® Rodent Diet 20, LabDiet, St. Louis, MO, USA). All mice were group-housed with up to 5 mice per ventilated cage, received water *ad libitum*, were kept on a 12:12 hr light:dark cycle at 22–23°C with 30–70% humidity, and had access to a small house and tissues for nesting. Adequate measures were taken to minimize pain and discomfort. Investigation has been conducted in accordance with the ethical standards and according to the Declaration of Helsinki and national and international guidelines and has been approved by the authors’ institutional review board. Mice were euthanized by 200 mg/kg sodium pentobarbital intraperitoneally injection followed by cervical dislocation. Blood was collected by cardiac puncture and major organs including liver, kidney, and spleen collected, rapidly frozen on dry ice and stored at −80 °C.

### COX Staining

Frozen liver tissues were embedded (Tissue-Tek; Sakura Finetek, Alphen aan den Rijn, The Netherlands), and 14-μm cryostat (Microm Model HM 500M Cryostat; Microm, Thermo Scientific, Fremont, CA, USA) sections taken at −21 °C were thawed onto slides (Super Frost; Menzel-Gläser, Thermo Scientific, Fremont, CA, USA) and stored at −80 °C until use. To visualize respiratory dysfunction, we used enzyme histochemistry to determine the activities of cytochrome c oxidase (COX) as previously described (24). Briefly, frozen sections were air dried for 1 h. Sections were then incubated for 40 min at 37 °C with 4 mM 3,3′-diaminobenzidine tetrahydrochloride (Sigma Liquid Substrate System D7304; Sigma-Aldrich, Sanit Louis, MO, USA), 100 μM cytochrome c, and 20μg/mL bovine catalase in 0.1M PBS pH 7. The sections were then washed four times at 10 min in 0.1 M PBS (pH 7.0), dehydrated in increasing concentrations of ethanol (70%, 95%, and 99.5%), coverslipped using Entellan^TM^ hard mounting medium (Sigma-Aldrich) and visualized under brightfield microscopy (Zeiss Axiophot 2 and Axioplan 2 Imaging Software, Zeiss Axiophot 2; Carl Zeiss, Oberkochen, Germany). Images were processed and quantified using appropriate software (FIJI v2.1.0/1.53c, Madison, WI, USA) (25).

### Oil Red O Staining

Frozen liver, heart, and skeletal muscle tissues were embedded (Tissue-Tek), and 14-μm cryostat (Microm Model HM 500M Cryostat) sections taken at −21 °C and stored at −80 °C until use. To visualize the presence of lipid droplets and fats in liver samples, Oil Red O staining was used as previously described (26,27). Tissue sections were air dried for 30 min and then immersed in 10% neutral buffered formalin for 10 min. Sections were then dipped in 60% isopropanol and stained in the working Oil Red O solution (Oil Red O, #00625; Sigma-Aldrich) for 15 min (the working Oil Red O solution was prepared and left to rest for 10 min). Sections were then again dipped in 60% isopropanol followed by deionized water and were finally coverslipped with an aqueous mounting medium (VectaMount, #H-5501; Vector Laboratories, Newark, CA, USA). Sections were visualized under brightfield microscopy (Zeiss Axiophot 2 and Axioplan 2 Imaging Software, Zeiss Axiophot 2; Carl Zeiss, Oberkochen, Germany), and images were processed and quantified using appropriate software (FIJI v2.1.0/1.53c, Madison, WI, USA) (25).

### Western Blot

Western blot was performed as previously described with minor changes (28). Briefly, tissues were lysed in radioimmune precipitation buffer (150 mm Tris-HCl, pH 7.5, 150 mm NaCl, 2 mm EDTA, 1% Triton, 0.1% SDS, and protease inhibitors), and protein concentration was measured using a BCA protein assay kit (Bio-Rad, Hercules, CA, USA). Fifty micrograms of total proteins were fractionated using NUPAGE® Novex® Bis-Tris precast gels (Invitrogen, Waltham, MA, USA) and blotted on 0.2-μm nitrocellulose membranes (Amersham Biosciences Hybond ECL, GE Healthcare, Chicago, IL, USA) as previously described. After incubation with the indicated primary and secondary antibodies, immunocomplexes were detected by chemiluminescence (PIERCE® ECL). Images were processed and quantified using appropriate software (FIJI v2.1.0/1.53c, Madison, WI, USA) (25). Antibodies used in this study are total OXPHOS Rodent WB Antibody Cocktail (ab110413), anti-CYP7A1 (ab65596), anti-POLGA (ab128899), anti-SREBP1 (ab28481) obtained from Abcam (Waltham, MA USA), anti-STAT1 (#9172), anti-STAT3 (#12640), anti-NF-kBp65 (#8242) obtained from Cell Signaling (Danvers, MA, USA), anti-GAPDH (sc-25778) obtained from Santa Cruz Biotechnology, Inc. (Dallas, Texas, USA), and anti-β-Actin−Peroxidase (A3854) obtained from MilliporeSigma (Burlington, MA, USA).

### Lipid Measurements

Sterols were measured in serum samples using gas chromatography mass spectrometry (GC-MS) after alkaline hydrolysis, as done previously (29), which allows for the quantitative analysis of cholesterol and structurally similar cholesterol precursors and plant sterols in a single chromatographic run. Briefly, pooled samples for cholesterol analysis were run on an Agilent HP 6890N gas chromatograph–Agilent HP 5973 MSD quadropole mass spectrometer using electron impact ionization mode (Stockholm, Sweden). Separation was performed on a HP-ultra1 (Scantec Lab, Gothenburg, Sweden) 25 m capillary column (0.20 mm i.d., 0.33 μm phase thickness). The samples for sterol analysis were run on an Agilent HP 5890 gas chromatograph– Agilent HP 5972 MSD quadropole mass spectrometer using electron impact ionization mode (Stockholm, Sweden), and separation was performed on a HP-5MS (Scantec Lab AB, Gothenburg, Sweden) 30 m capillary column (0.25 mm i.d., 0.25 μm phase thickness). Deuterium labelled cholesterol, sitosterol, and lathosterol were used as internal standards.

### Real Time PCR

Tissue samples (30–50 mg) were lysed, and RNA was extracted and purified (Zymo Direct-zol RNA MiniPrep Plus Kit R2070, Zymo Research, Irvine, CA, USA) according to the manufacturer’s instructions. RNA concentration was determined using a spectrophotometer (NanoDrop, ND-2000), and reverse transcription was run (Lunascript^®^ RT Supermix Kit E3010, New England BioLabs Inc., Ipswich, MA, USA) according to the manufacturer’s protocol. Once the cDNA was synthesized, qPCR reactions were run using a SYBR green-based master mix (Luna^®^ qPCR Mastermix M3003, New England BioLabs Inc.) according to the manufacturer’s instructions using appropriate instrumentation (Viia7 Real-Time PCR System, Applied Biosystems, Waltham, MA, USA). Results were analyzed using the appropriate software (GraphPad Prism v. 9, San Diego, CA, USA). The primer sequences used in this study are as follows: HMGCR_For_: CCGGCAACAACAAGATCTGTG, HMGCR_Rev_: ATGTACAGGATGGCGATG CA; HMGCS1_For_: CTCTGTCTATGGTTCCCTGGCT, HMGCS1_Rev_: TCCAATCCTCTTCCCTGCC (30); LDL_For_: AGGCTGTGGGCTCCATAGG, LDL_Rev_: GCGGTCCAGGGTCATCT (31); s100a8_For_: CCTTTGTCAGCTCCGTCTTCA, s100a8_Rev_: TCCAGTTCAGACGGCATTGT; s100a9_For_: AATGGTGGAAGCACAGTTGG, s100a9_Rev_: CTGGTTTGTGTCCAGGTCCTC; RNaseL_For_: CCTGGGATAAAGTCAGATGGA, RNaseL_Rev_: GGGATTGGGCTGATTATGTT; Usp18_For_: CGTCCTCTTCTGTAGCCCTT, Usp18_Rev_: ACAAATGACCCTCTCAAGCA; Isg15_For_: CTAGAGCTAGAGCCTGCAG, Isg15_Rev_: AGTTAGTCACGGACACCAG; Ifit1_For_: CAAGGCAGGTTTCTGAGGAG, Ifit1_Rev_: GACCTGGTCACCATCAGCAT; Ifit4_For_: TTCCCAGCAGCACAGAAAC, Ifit4_Rev_: AAATTCCAGGTGAAATGGCA; LTF_For_: AATCCAATCTCTGTGCCCTG, LTF_Rev_: ATGCAACATTTCCTGCCTTC; C1qa_For_: TGGACAGTGGCTGAAGATGT, C1qa_Rev_: AAACCTCGGATACCAGTCCG; Actb_For_: ACCTTCTACAATGAGCTGCG, Actb_Rev_: CTGGATGGCTACGTACATGG.

### Proteomic Analysis

We performed quantitative proteomics as previously described (32). Very briefly, liver tissues were lysed in SDS-lysis buffer, protein concentration was estimated using a DC protein assay kit (Bio-Rad, Hercules, CA, USA), and 250 μg of total protein from each sample was processed according to a modified FASP protocol (33). 100 μg of peptides from each sample were labeled with TMT10plex (Thermo Scientific) according to the manufacturer’s instructions. TMT-pooled sample sets then underwent high-resolution isoelectric focusing (HiRIEF) separation. Each HiRIEF fraction was then analyzed using liquid chromatography-mass spectrometry (LC-MS). All MS/MS spectra were searched by MSGF+/Percolator under the Galaxy platform (https://usegalaxy.org) using a target decoy strategy. The reference database used was the UniProt reference Mus musculus proteome (canonical and isoform, 51,529 protein entries, downloaded from uniprot.org on 2014 08 01). A precursor mass tolerance of 10 ppm and high resolution setting on MS2 level as used and only peptides with fully tryptic termini were allowed. Quantification of TMT 10plex reporter ions was done using an integration window tolerance of 10 ppm. PSMs and peptides were filtered at 1% FDR (peptide level), and proteins were filtered additionally at 1% FDR (protein level) using the “picked” protein FDR method (34). The mass spectrometry proteomics data have been deposited to the ProteomeXchange Consortium via the PRIDE partner repository with the dataset identifier PXD047574. To compare protein levels between conditions, Student’s t-tests were applied to compare protein levels between WT and mtDNA mutator mice on log2 transformed data using SAM (Significance Analysis of Microarrays, http://statweb.stanford.edu/~tibs/SAM/) as previously done for LC MS/MS data (32). Significantly de regulated proteins (FDR ≤ 0.05) were analyzed with the freely accessible WEB based software analysis tool GEne SeTAnaLysis Toolkit (WebGestalt) (http://www.webgestalt.org/option.php) (35). Gene Ontology (GO) enrichment analysis for biological processes, molecular function, and cellular components (36), Kyoto Encyclopedia of Genes and Genomes (KEGG) for pathway enrichment analysis in disease (37).

### Cell Culture

Primary fibroblast cell cultures were prepared as previously reported (38). Briefly, tail clips from Mut and WT mice were placed in 10 cm Petri dishes, washed three times with phosphate buffer saline (PBS), and diced into small pieces with a razor blade. Tissue pieces were collected in 15 mL Falcon tubes and spun down at 1000 g for 5 min. After removing the PBS, 1 mL of a 0.25% trypsin-EDTA solution was added to the tissue, followed by 37 °C incubation for 30 min with 350 rpm shaking. After digestion, cells were collected by 1000 *g* centrifugation for 5 min, and the 0.25% trypsin-EDTA solution was replaced by IMDM containing L-glutamine supplemented with 20% FBS, 100 IU/mL penicillin, and 50 μg/mL streptomycin. Cells were plated in a 100 mm Petri dish and grown in a 37 °C, 5% CO_2_ humidified incubator. After reaching 70% confluence, cells were split 1:3 to 1:4 into T75 flasks and the IMDM replaced with DMEM with the percentage of FBS lowered to 10%. To assess the mitochondrial membrane potential (MMP), cells were incubated for 30 min in DMEM containing 100 nM tetramethylrhodamine (TMRM) (Thermo Fisher Scientific, Waltham, MA). After 30 min, cells were washed three times with PBS, collected by trypsinization, and the TMRM fluorescence intensity measured by flow cytometry (Ex/Em: 488 nm/570 nm). To determine the effect of the oligomycin inhibitor on MMP, cells were incubated for 24 h with oligomycin (Sigma-Aldrich) at concentrations indicated in the figure. The level of intracellular ROS was assessed using the CM-H2DCFDA indicator (Thermo Fisher Scientific). Cells were incubated 1 h with DMEM containing either 10 μM CM-H2DCFDA alone or in the presence of 100 μM H_2_O_2_. Thereafter, the indicator containing media was replaced with regular DMEM and fluorescence intensity was measured by flow cytometry (Ex=488nm). Cellular oxidative stress was assessed by measuring the conversion rate of the oxidative-sensitive fluorescent protein mitoTIMER (Addgene plasmid #50547) (39). The encoding sequence for mitoTIMER was subcloned into the Tet-On pTRIPZ lentivirus backbone (Dharmacon™ Reagents, GE Healthcare, Lafayette, CO, USA). Lentivirus was generated as previously described, and fibroblasts were transduced accordingly (40). Four days post-transduction, cells were selected in DMEM containing 1 µM puromycin. For the experiment, cells were seeded in 60 mm Petri dishes, and mitoTIMER expression was induced for either 24 or 48 h by adding 0.1 µg/mL doxycycline to the media. MitoTIMER fluorescence was measured by flow cytometry using both the FITC and TRITC channels (Ex = 488 nm and 550 nm), and the mitoTIMER conversion rate expressed as the Red/Green ratio. To generate Rho0 cells, mtDNA mutator-derived fibroblasts were infected with a lentivirus expressing Cre recombinase and mCherry fluorescence protein. Lentiviral plasmid pLM-CMV-R-Cre was obtained from Addgene (#27546) (41). After infection, cells were sorted for mCherry expression, and the media was supplemented with 50 μg ml^-1^ uridine.

### mtDNA Copy Number

Relative mtDNA copy number was determined as previously done (32). Briefly, after total DNA extractions were prepared, real time PCR was performed using two different sets of primers were directed to the mitochondrial genes mtCO2 and mtCO1 and the nuclear genes RSP18 and NDUFV1, respectively. Primer sequences were as follows: mtCO2_fw_ ATAACCGAGTCGTTCTGCCAAT and mtCO2_rev_ TTTCAGAGCATTGGCCATAGAA; RSP18_fw_ TGTGTTAGGGGACTGGTGGACA and RSP18_rev_ CATCACCCACTTACCCCCAAAA; mtCO1_fw_ TGCTAGCCGCAGGCATTAC and mtCO1_rev_ GGGTGCCCAAAGAATCAGAAC; and NDUFV1_fw_ CTTCCCCACTGGCCTCAAG and NDUFV1_rev_ CCAAAACCCAGTGATCCAGC. Relative fold changes were calculated with the delta Ct method for each set of primers and averaged. Data were normalized against WT levels.

### Mitochondrial Respiration

To assess mitochondrial respiration in WT- and mtDNA mutator-derived primary fibroblasts, we performed a Mito Stress Test using a Seahorse XFe96 Analyzer (Agilent, Santa Clara, CA). 50,000 cells were seeded into Seahorse XF96 tissue culture microplates and incubated at 37 °C overnight. Cell culture media was replaced 1 h before analysis with Seahorse XF DMEM Medium, pH 7.4 (103575-100) supplemented with 10 mM glucose, 1mM pyruvate, and 2mM L-Glutamine, and drugs were loaded into the Seahorse XFe96 Sensor Cartridge (Final concentrations: Oligomycin 1 µM, FCCP 1 µM, Antimycin/Rotenone ‘R/A’ 1 µM) according to the manufacturer’s specifications. Seahorse analysis was performed at 37 °C. Mitochondrial respiration recordings were normalized to cell number using CyQUANT (Thermo Fisher C7026) fluorescence on a plate reader. Data analysis was performed using Seahorse Wave Desktop Software 2.6, Excel, and Prism. N=6 wells for each condition. Significance was calculated using an unpaired t-test.

### Immunocytochemistry

Immunocytochemistry was performed as previously described (42). Briefly, cells grown on glass cover slides were rinsed three times with PBS and fixed with 4% formaldehyde in PBS for 15 min. After fixation, cells were permeabilized with 0.1% Triton X-100 and 4% bovine serum albumin in PBS for 30 min. Incubation with the indicated primary antibody was conducted for 1 h. Thereafter, cells were rinsed with PBS and incubated for 1 h with the appropriate fluorochrome conjugated secondary antibody (GenTex, Montréal, Canada) and washed three times with PBS before mounting with VECTASHIELD medium containing 4′,6-Diamidino-2-Phenylindole Dihydrochloride (DAPI) (Vector Laboratories Inc. Burlingome, CA, USA). Immunofluorescence was examined using a confocal microscope (Olympus Fluoview FV1000) or with a wide-field fluorescence microscope (Nikon ECLIPSE T*i* and Nikon Elements Imaging Software v4.13, Tokyo, Japan). Images were processed and quantified using appropriate software (FIJI v2.1.0/1.53c, Madison, WI, USA) (43).

### Statistical Analysis

Data are presented to three significant digits as mean values ± SEM with sample size (N) indicated in the figure legends. The statistical analyses were performed using appropriate software (GraphPad Prism v. 9, San Diego, CA, USA) and include: unpaired t-test, one-way ANOVA, and two-way ANOVA with Tukey post-hoc multiple comparisons, with an α level of 0.05. Significances are denoted in figures with * *p* < 0.05, ** *p* < 0.01, *** *p* < 0.001, **** *p* < 0.0001, and α as trending with *p* < 0.10.

## Results

### Liver Steatosis in mtDNA Mutator Mice

The phenotype of mtDNA mutator (Mut) mice revealed an enlarged liver apparent at 20-25 weeks, progressing until mortality within the first year, around 50 weeks of age (Fig. 1A). At 25 weeks, the Mut liver weight, normalized to body weight, was approximately 20 % larger than wild-type (WT) controls, increasing to over 30 % at 45 weeks of age (Fig. 1B). Notably, normalized liver weights increased as well in WT mice with age. To determine the impact of mtDNA mutations on OXPHOS Complexes, key subunits levels were assessed by Western blot. Cytochrome *c* oxidase subunit 1 (MTCO1) in Complex IV displayed significant down-regulation in Mut liver extracts at both younger (25-30 weeks) and older (40-45 weeks) ages, resembling the pattern observed in older WT mice (>110 weeks), while the nuclear encoded succinate dehydrogenase B (SDHB) in Complex II was up-regulated in younger and older Mut mice as compared to WT (Fig. 1C and D). COX staining further demonstrated a substantial decrease in complex IV activity in Mut liver compared to WT controls (Fig. 1E). To determine if the observed increase in liver size could be due to intracellular accumulation of lipid droplets, Oil Red O staining was used to label lipid droplets and revealed a marked intracellular lipid accumulation in Mut liver as compared to WT, starting already at 25 weeks of age and surpassing that observed in normal aging, while no such accumulation was found in heart and skeletal muscle tissues (Fig. 1F), indicating that accumulation of lipid droplets secondary to mitochondrial dysfunction is specific to liver tissue.

**Figure 1.**
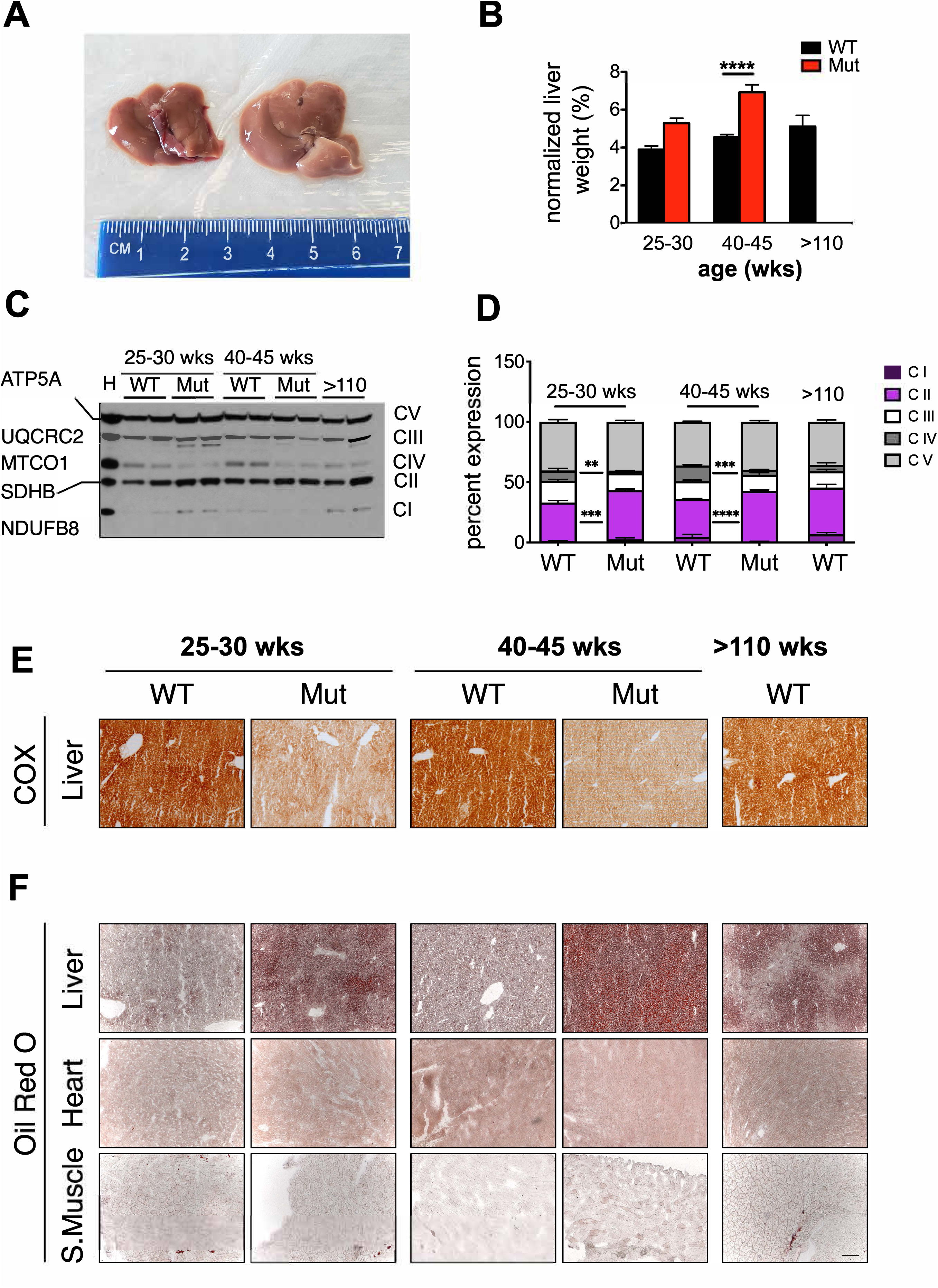
Mitochondrial dysfunction induces fatty liver. **(A)** Representative images of wild-type (WT) and mtDNA mutator mouse (Mut) livers from ≅ 42 wks old males. **(B)** Liver weights normalized by body weights in 25-30 week-old mtDNA mutator (Mut, red, N=4) and wild-type (WT, black, N=4) mice, 40-45 week-old mtDNA mutator (Mut, red, N=14) and wild-type (WT, black, N=21) mice, and >110 week-old WT (black, N=5) mice. Significances were determined by two-way ANOVA and Sidak’s multiple comparison test. Data are presented as mean ± SEM with **** p < 0.0001. **(B) (C)** Representative Western blot with a batch of 5 antibodies, each recognizing a protein belonging to one of oxidative phosphorylation (OXPHOS) complexes. Proteins (ATP5A, UQCRC2, MTCO1, SDHB, and NDUFB8) are presented on the left-hand side while Complexes (CI, CII, CIV, CIII, CV) are presented on the right. Heart lysate (H) was used for reference in the first column. **(D)** Western blot quantification showing the percent expression of CI (dark purple), CII (light purple), CIII (white), CIV (dark grey), and CV (light grey) complexes individually. Mut mice (N=4 per age) and WT mice (N=4 per age) were analyzed at weeks 25-30, 40-45, and >110 (WT only). Significances were determined by two-way ANOVA and Tukey’s post-hoc analysis with **p < 0.01, ***p < 0.001, and ****p< 0.0001. **(E)** Representative COX staining of liver sections from Mut and WT mice aged 25-30 weeks, 40-45 weeks, and older than 110 weeks (WT only). **(F)** Representative Oil Red O staining of liver, heart, and skeletal muscle sections from Mut and WT mice aged 25-30 weeks, 40-45 weeks, and older than 110 weeks (WT only). Scale bar: 200 µm.

### Lipid Metabolism in mtDNA Mutator Mice

To evaluate the impact of liver steatosis on lipid metabolism, serum cholesterol and triglyceride levels were measured in WT and Mut mice pooled samples at different ages. While Mut mice exhibited lower serum cholesterol levels at younger ages, a reversal occurred at 43-45 weeks, with Mut mice displaying higher cholesterol levels than WT mice (Fig. 2A). In contrast, despite increasing with age, serum triglyceride levels in Mut mice remained consistently lower than those in WT mice across all analyzed ages (Fig. 2B). Analysis of serum lathosterol and 7α-hydroxy-4-cholesten-3-one (C4) levels indicated a potential impairment in cholesterol secretion via bile acid synthesis in Mut mice, particularly at older ages rather than an increase in cholesterol synthesis (Fig. 2C and D). Since bile acid is also needed to metabolize phytosterols, which are plant sterols similar to cholesterol, we measured the serum level of two phytosterols: sitosterol and campesterol and found that the serum level of both were elevated in Mut mice at older age, further supporting a possible impairment in their secretion via bile acid synthesis (Fig. 2E and F).

**Figure 2.**
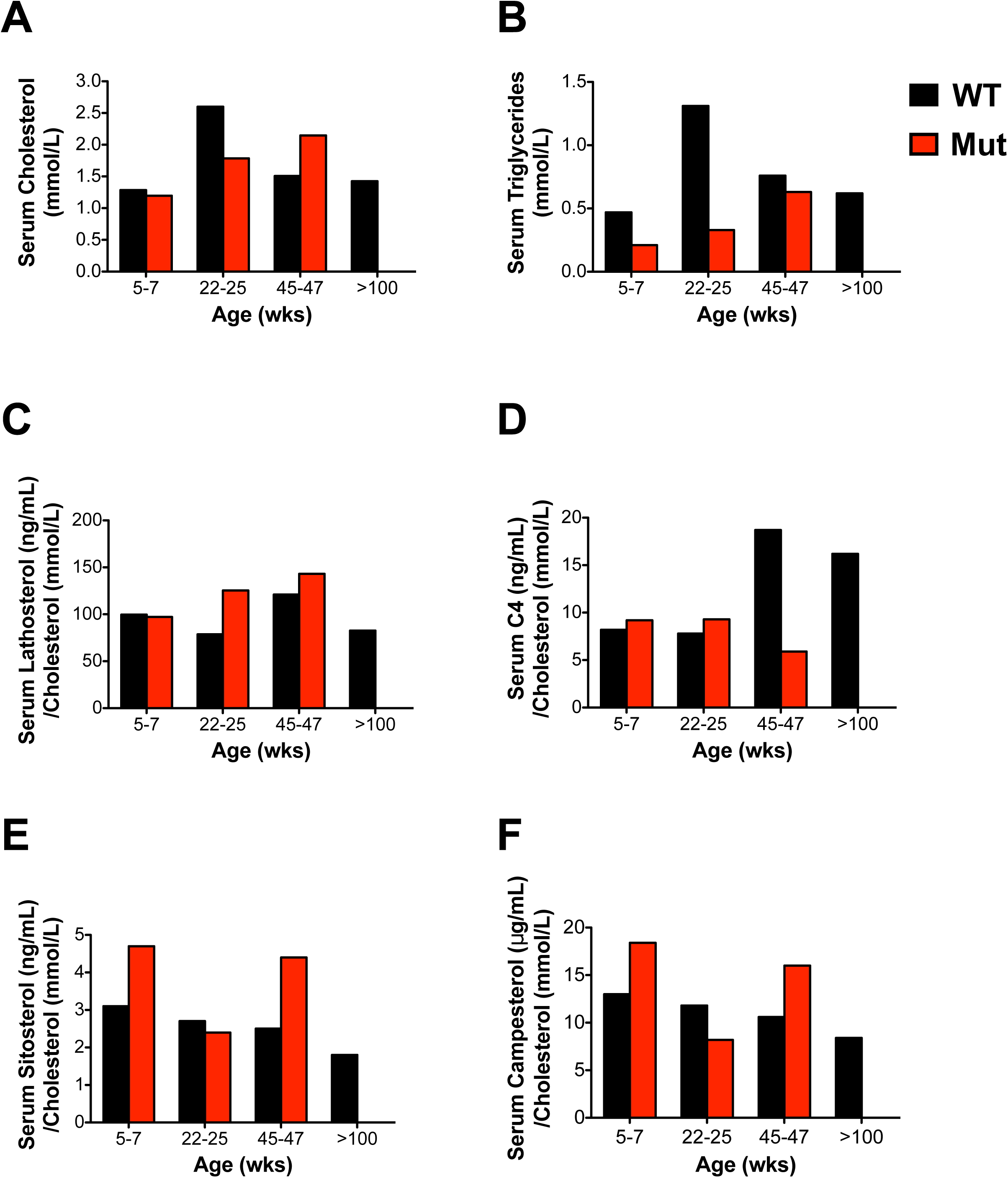
Mitochondrial dysfunction affects cholesterol levels in serum. HPLC measurement of serum blood level of: **(A)** cholesterol (mmol/L), **(B)** triglycerides (mmol/L), **(C)** ratio of lathosterol (ng/mL) to cholesterol (mmol/L), **(D)** ratio of C4 (ng/mL) to cholesterol (mmol/L), **(E)** ratio of sitosterol (ng/mL) to cholesterol (mmol/L), **(F)** ratio of campesterol (ng/mL) to cholesterol (mmol/L), in mtDNA mutator (Mut, red) and wild-type (WT, black) mice aged 5-7 weeks, 22-25 weeks, 45-47 weeks, and WT >100 weeks. Each column represents N=9-11 mice pooled together.

### Analysis of Liver Proteins in Mut Mice

Protein analysis in Mut liver revealed surprising findings. CYP7A1, a key enzyme in the cholesterol catabolic pathway, which converts cholesterol to bile acids, exhibited higher protein levels when assessed by Western blot in Mut mice at both younger and older ages, suggesting a tendency to upregulate the bile acid synthesis pathway (Fig. 3A and B). Intriguingly, the sterol regulatory element-binding protein 1 (SREBP-1), a master regulator of cholesterol biosynthesis, showed a dramatic increase in its precursor and cleaved active forms in older Mut mice (Fig. 3C), indicating a complex interplay in lipid synthesis regulation. Such up-regulation was specific for liver tissue as it was not found in kidney and skeletal muscle tissues (Fig. 3D and E). Further investigation revealed a downregulation of 3-hydroxy-3-methylglutaryl-CoA reductase (HMGCR), the rate-limiting enzyme of cholesterol synthesis, and low-density lipoprotein receptor gene (LDLR) with age in Mut mice, while expression of 3-hydroxy-3-methylglutaryl-CoA synthase 1 (HMGCS1), a gene involved in cholesterol synthesis (HMGCS1), exhibited varying trends (Fig. 3F, G, and H). Despite SREBP-1 activation, genes involved in cholesterol uptake and synthesis were paradoxically down-regulated as liver steatosis progressed.

**Figure 3.**
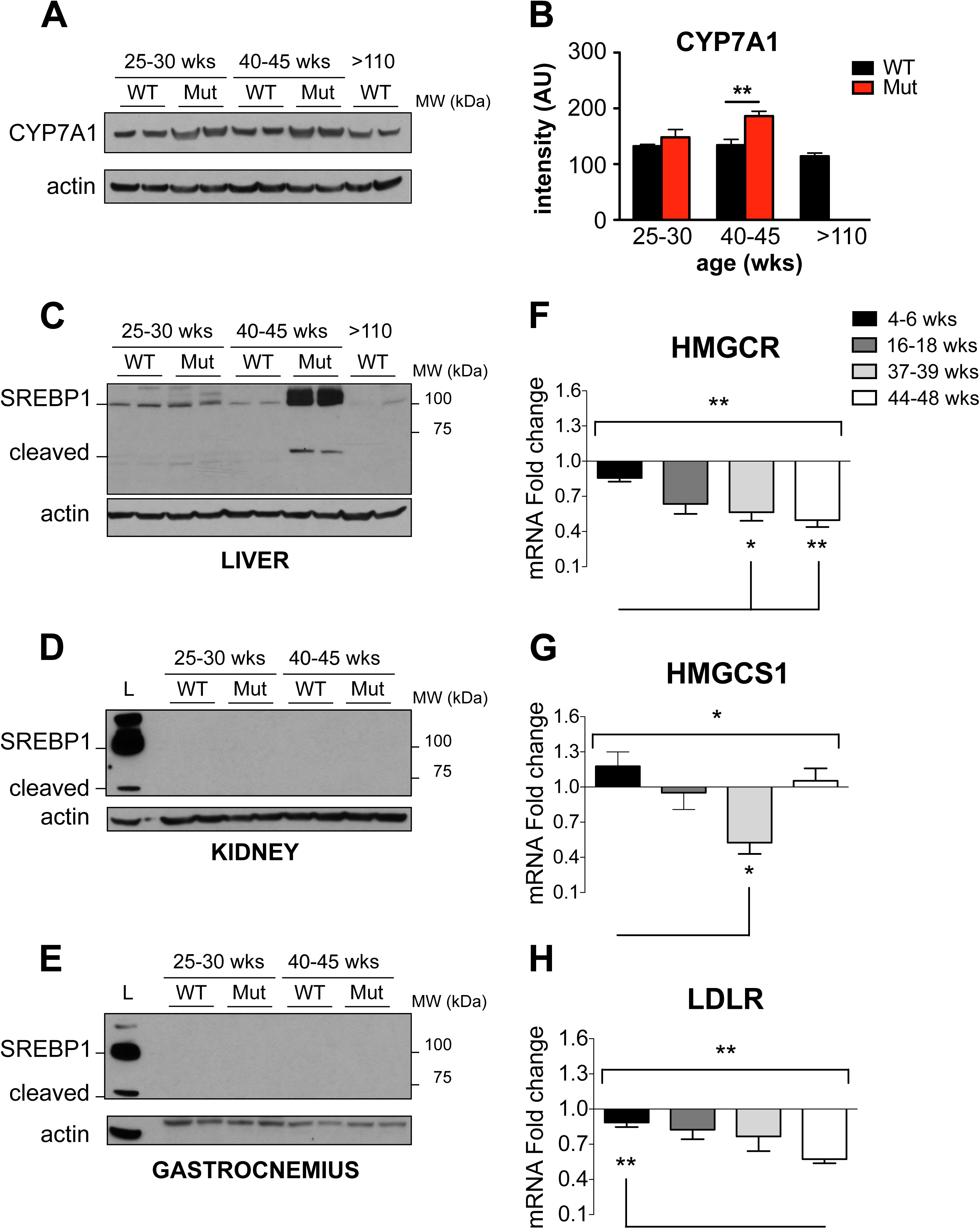
Mitochondrial dysfunction alters expression of proteins involved in cholesterol metabolism in liver. **(A)** Representative Western blot showing CYP7A1 levels in liver from mtDNA mutator (Mut) and wild-type (WT) mice analyzed at 25-30, 40-45, and >110 weeks (WT only). **(B)** CYP7A1 western blot quantification of Mut (red, N=4 per age) and WT (black, N=4 per age) mice aged 25-30, 40-45, and >110 (WT only) weeks. Significances were determined by two-way ANOVA and Tukey’s post hoc analysis with **p < 0.01. **(C)** Representative western blot showing total and activated (cleaved) transcription factor SREBP1 in liver from 25-30 and 40-45 week-old Mut and WT mice as well as from more than 110 week-old WT mice. **(D-E)** SREBP1 transcription factor immunoblot of kidney (D) and gastrocnemius (E) from Mut (N=4 per age) and WT (N=4 per age) mice aged 25-30 and 40-45 weeks. First column showing 45 wk-old Mut liver sample (L) for reference. **(F, G, H)** qPCR measuring mRNA fold-change of HMGCR, HMGCS1, and LDL receptor in liver from Mut as compared to WT mice aged 4-6 weeks (black, N=6 each group), 16-18 weeks (dark grey, N=4 each group), 37-39 weeks (light grey, N=4 each group), and 44-48 weeks (white, N=6 each group). Significances were determined by one-way ANOVA and Bonferroni’s post-analysis test with *p < 0.05, and **p < 0.01.

To gain deeper insights into the diverse biological processes affected by liver steatosis induced by mitochondrial dysfunction, a quantitative proteomic analysis comparing WT and Mut male mice liver proteins was conducted. Of the over 8,000 identified proteins, 604 exhibited significant (FDR=0.0) differential regulation, with 346 upregulated and 258 downregulated. Gene ontology analysis indicated an association of up-regulated proteins with inflammatory processes, while down-regulated proteins were primarily linked to mitochondrial processes such as the OXPHOS pathway (Fig. 4A and B). The analysis also revealed significant associations with coenzyme Q reductase deficiency, liver cirrhosis, and non-alcoholic fatty liver disease (NAFLD) (Fig. 4C). Focusing on the 69 proteins associated with NAFLD, 56 were up-regulated and 13 down-regulated, emphasizing the profound impact of mitochondrial dysfunction on liver pathology (Fig. 4D).

**Figure 4.**
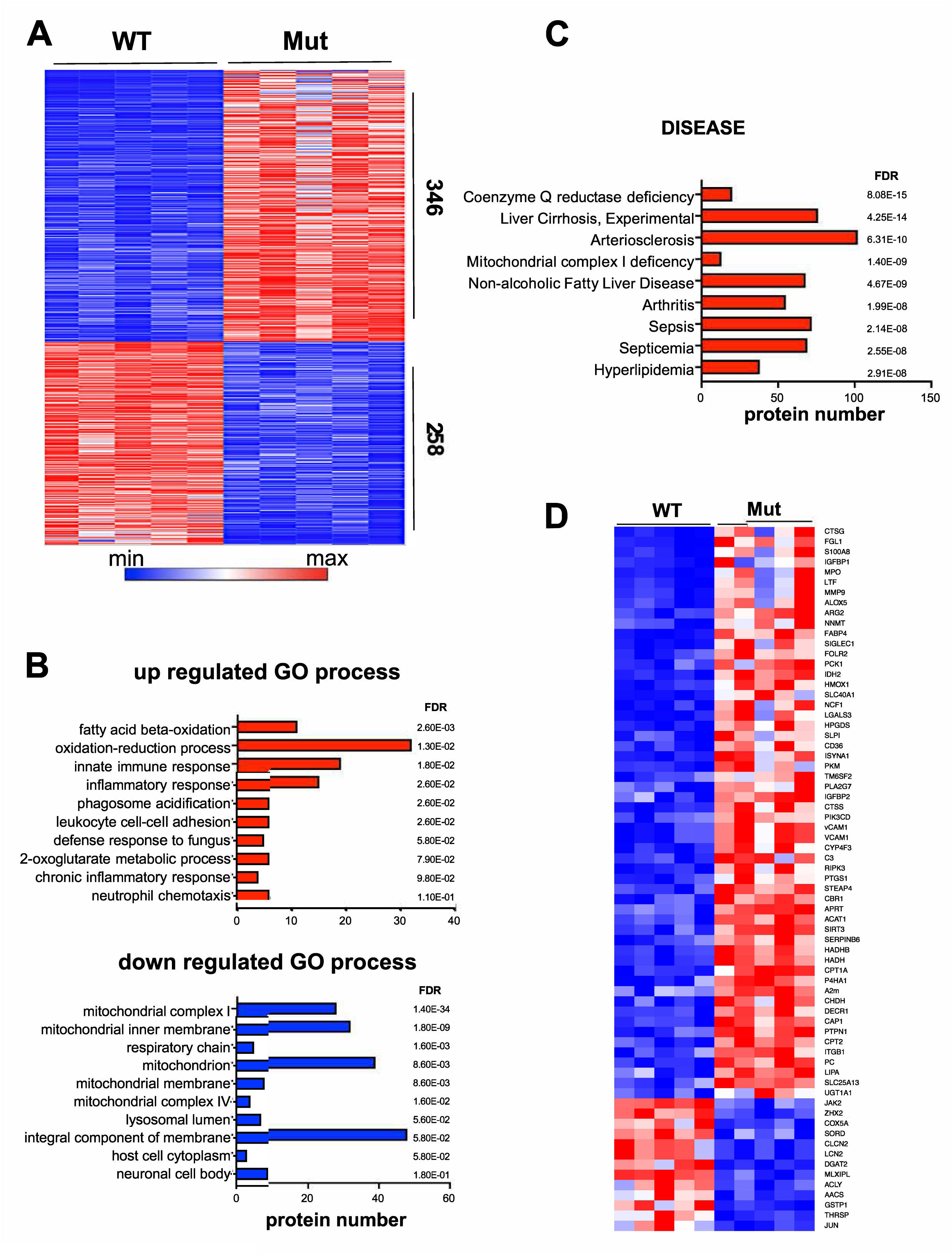
Proteomic analysis of liver protein. **(A)** Heat map representing proteomic liver analysis from 45±2 week-old mtDNA mutator mice (Mut, N=5) as compared to age-matched wild-type (WT, N=5) mice. 346 proteins were found to be significantly up-regulated (red) and 258 proteins were down-regulated (blue) in the Mut mice. The false discovery rate (FDR) for all 604 proteins shown is 0. **(B)** Gene ontology (GO) analysis of up- (red) and down- (blue) regulated biological processes in mtDNA mutator mice. **(C)** Identified diseases associated with the proteomic profile of mtDNA mutator mice with up- and down-regulated proteins analyzed together. **(D)** Heat map showing individual proteins (69) associated with non-alcoholic fatty liver disease (NAFLD) in mtDNA mutator mice as compared to WT littermates. The majority of proteins (56) were significantly up-regulated (red) while a few (13) proteins were down-regulated (blue) in Mut mice.

### Mitochondrial Dysfunction and Inflammation

One surprising outcome of our proteomic analysis was the identification of numerous proteins associated with inflammation showing upregulation in the liver tissue from mtDNA mutator mice (Fig. 5A). This suggests that mtDNA mutations induce steatosis and inflammation in the liver. Notably, several upregulated proteins were genes induced by interferon type I activation. To validate our proteomic data and determine whether these protein changes were due to increased gene expression, we chose to assess the mRNA expression of 9 genes identified as up-regulated (S1008A, S1009A, LTF, C1qA, Ifit1, Ifit4, RNAseL, USP18, and ISG15) in our analysis of liver from Mut and WT mice using real-time PCR analysis. Our findings indicate that the expression of 7 out of 9 genes were significantly up-regulated in liver samples from mtDNA mutator mice, as compared to WT littermates, thus confirming our proteomic data (Fig. 5B). Since the mtDNA mutator mouse is a model of progeria and mitochondrial dysfunction is a hallmark of aging, we also assessed the expression of these 9 genes in liver samples from young (3 months old) and old (24 months old) healthy C57BL/6J mice. We found that 4 out of the 9 genes were significantly up-regulated in old mice compared to young controls, suggesting an ongoing inflammatory process. However, the level of up-regulation was much less compared to what was observed in liver samples from mtDNA mutator mice (Fig. 5C).

**Figure 5.**
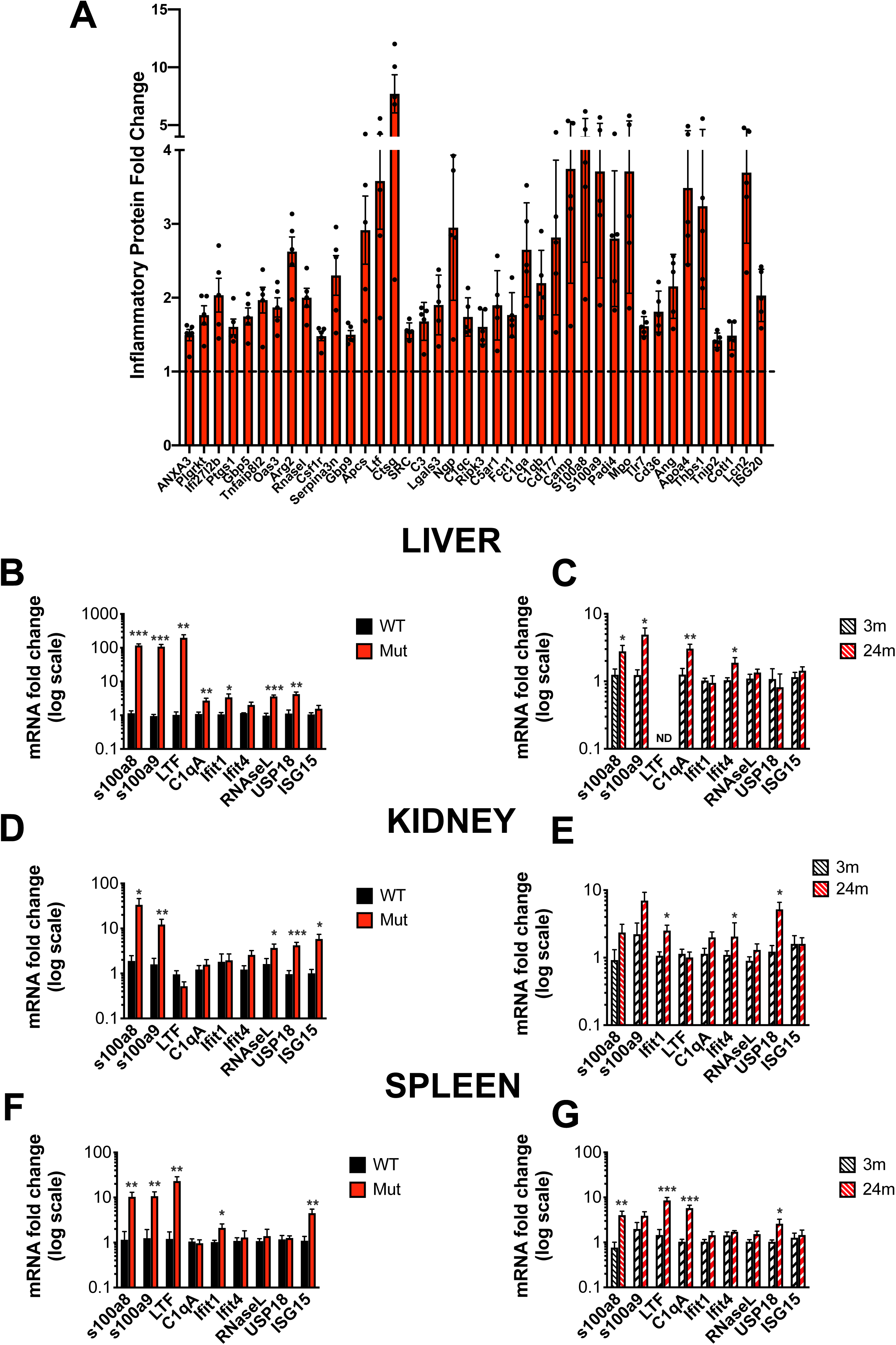
Mitochondrial dysfunction triggers inflammation. **(A)** Inflammatory proteins significantly up-regulated in liver from 45±2 week-old mtDNA mutator mice compared to age-matched WT littermates, as determined by the proteomic analysis (N=5 per group). **(B-G)** mRNA fold change in inflammatory genes assessed by qPCR in liver **(B, C)**, kidney **(D, E),** and spleen **(F, G)** from ∼45 week-old Mut (red, N∼9) as compared to WT (black, N∼9) mice as well as 3 month-old (black-stripe, N∼9) and 24 month-old (red-stripe, N∼9) C57B/6J WT mice. Changes are reported on a logarithmic scale. Significant differences were assessed by unpaired t-test with *p < 0.05, **p < 0.01, and ***p < 0.001.

To determine whether the up-regulation of inflammatory genes is restricted to the mtDNA mutator liver tissue or is a more widespread phenomenon affecting other tissues/organs, we assessed the expression of these 9 genes in the kidney and spleen from mtDNA mutator and WT mice. We found that several genes were also up-regulated in mtDNA mutator tissues compared to WT, suggesting that chronic inflammation underlies the mutator mouse phenotype (Fig. 5D and F). Analysis of kidney and spleen samples from young and old mice confirmed the up-regulation of inflammatory-associated genes in old tissues as well (Fig. 5E and G).

### Mitochondrial Dysfunction in mtDNA mutator-Derived Fibroblasts

To further investigate the metabolic and inflammatory changes induced by mitochondrial dysfunction in mtDNA mutator mice, we generated several fibroblast cell lines from Mut and WT mice. Quantitative PCR analysis of mtDNA content in these cell lines revealed that mtDNA mutator-derived cells have an overall lower number of mtDNA copies compared to WT cells (Fig. 6A). Surprisingly, analysis of their respiratory capacity using the Agilent Seahorse instrument demonstrated that mtDNA mutator-derived cells exhibit a significantly lower oxygen consumption rate (OCR), as compared to WT cells. This reduction in OCR is unresponsive to Complex V inhibition by oligomycin, to membrane potential dissipation by the potent OXPHOS uncoupler FCCP, or to electron transport chain (ETC) blockage by rotenone and antimycin A (Fig. 6B). Interestingly, inhibition of ETC by rotenone and antimycin A reduced the OCR of WT cells to the level observed in the mtDNA mutator-derived fibroblasts (Fig. 6B). Furthermore, basal respiration, ATP production, and spare respiratory capacity were all significantly lower in the mtDNA mutator-derived cells, as compared to WT cells (Fig. 6C, D, and E). In contrast, the extracellular acidification rate (ECAR), an indicator of glycolysis, was significantly higher in the mtDNA mutator fibroblasts compared to WT cells (Fig. 6F and G). Importantly, WT cells exhibited ECAR values similar to those of mtDNA mutator cells only when OXPHOS was inhibited by either oligomycin, rotenone, or antimycin A (Fig. 6F). Taken together, our data demonstrate that mtDNA mutator-derived cells rely heavily on glycolysis for ATP energy production.

**Figure 6.**
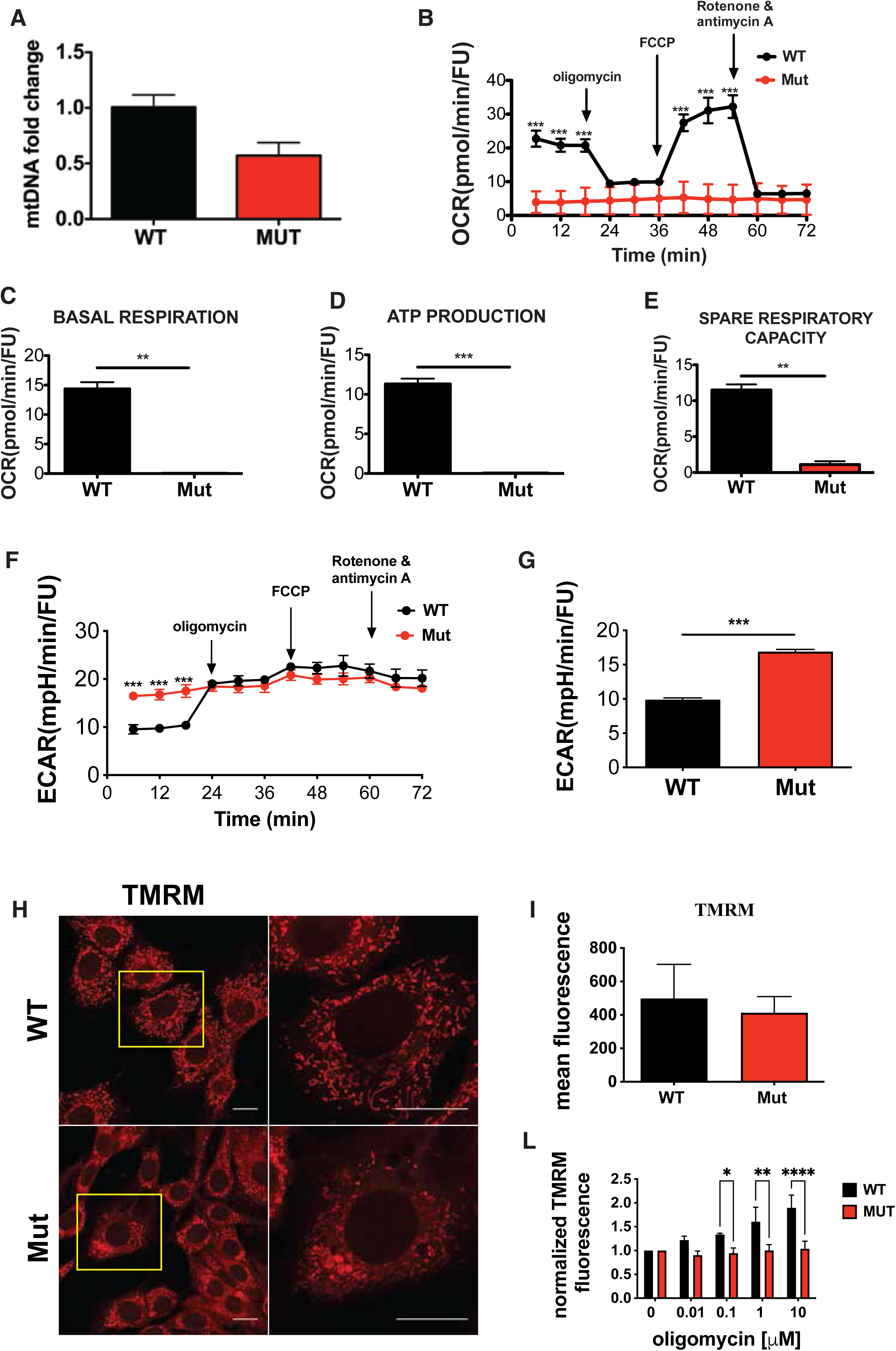
Mitochondrial function in primary-derived fibroblast cell lines. **(A)** mtDNA levels assessed by qPCR in two different fibroblast cell lines derived from mtDNA mutator (Mut) and wild-type (WT) mice. **(B)** Oxygen consumption rate (OCR) in Mut and WT-derived fibroblast cell lines as measured by Agilent Seahorse XF Analyzer. Each condition was tested with four replicates and significance was determined by two-way ANOVA with Tukey’s post-hoc analysis, ***p < 0.001. **(C, D, E)** Bar graphs representing basal oxygen consumption rate (OCR), ATP production, and spare respiratory capacity, respectively, based on OCR in Mut and WT-derived fibroblasts. Significance assessed by unpaired t-test with **p < 0.01 and ***p < 0.001. **(F and G)** Extracellular acidification rate (ECAR) in Mut and WT-derived fibroblasts under basal condition and in response to oligomycin, FCCP, and rotenone with antimycin A treatments. ECAR was measured using Agilent Seahorse XF Analyzer and significance was determined by unpaired t-test with ***p < 0.001. **(H)** Representative images of mitochondrial network stained with tetramethylrhodamine methyl ester (TMRM) in WT and Mut mice. Bar scale 20 µm. **(I)** Bar graph showing TMRM mean fluorescence determined by flow cytometry (FACS) in two different fibroblast derived cell lines from WT and Mut mice. **(J)** TMRM mean fluorescence determined by FACS in three mtDNA mutator (Mut) and two WT-derived fibroblast cell lines. Cells were treated with different concentrations of oligomycin for 24 h and data were expressed as percentage of the control. Significances were determined by two-way ANOVA and Bonferroni’s post-analysis test with *p < 0.05, **p < 0.01, and ***p < 0.001.

To investigate the effect of OXPHOS impairment on mitochondrial membrane potential (ΔΨm) in mtDNA mutator cells, we used tetramethylrhodamine methyl ester (TMRM) to assess ΔΨm in both mtDNA mutator- and WT-derived fibroblasts. Microscopic observation of the mitochondrial network in cells incubated with TMRM revealed that while WT cells exhibited a healthy mitochondrial network characterized by elongated thin mitochondria, mtDNA mutator cells displayed disrupted mitochondrial network organization with the accumulation of several swollen mitochondria in their cytoplasm (Fig. 6H). Surprisingly, no significant difference in ΔΨm between mtDNA mutator and WT cells was found when TMRM intensity was measured using flow cytometry analysis (Fig. 6I). However, when we blocked Complex V with different concentrations of oligomycin, WT cells exhibited a correlation between ΔΨm increase and oligomycin concentrations, whereas no change in ΔΨm was induced in mtDNA mutator cells upon treatment with oligomycin (Fig. 6H).

To understand if mitochondrial dysfunction in mtDNA mutator-derived fibroblasts induces oxidative stress, we examined the levels of reactive oxygen species (ROS) using the CM-H2DCFDA colorimetric probe. While no significant differences in oxidative stress were observed when comparing mtDNA mutator cells to controls, Mut cells displayed reduced antioxidant capability upon H_2_O_2_ treatment, resulting in significantly higher levels of oxidative stress (Fig. 7A and B). To further investigate the level of oxidative stress in mtDNA mutator fibroblasts, we utilized the mTIMER protein. The green-to-red conversion of mTIMER, reflecting chromophore oxidation, was monitored in mtDNA mutator and WT cells stably transduced with a lentivirus expressing mTIMER under a dox-regulated promoter. The results showed that the green-to-red conversion was significantly faster in mtDNA mutator cells compared to WT, further suggesting that mtDNA mutator cells have a higher oxidative environment (Fig. 7C and D).

**Figure 7.**
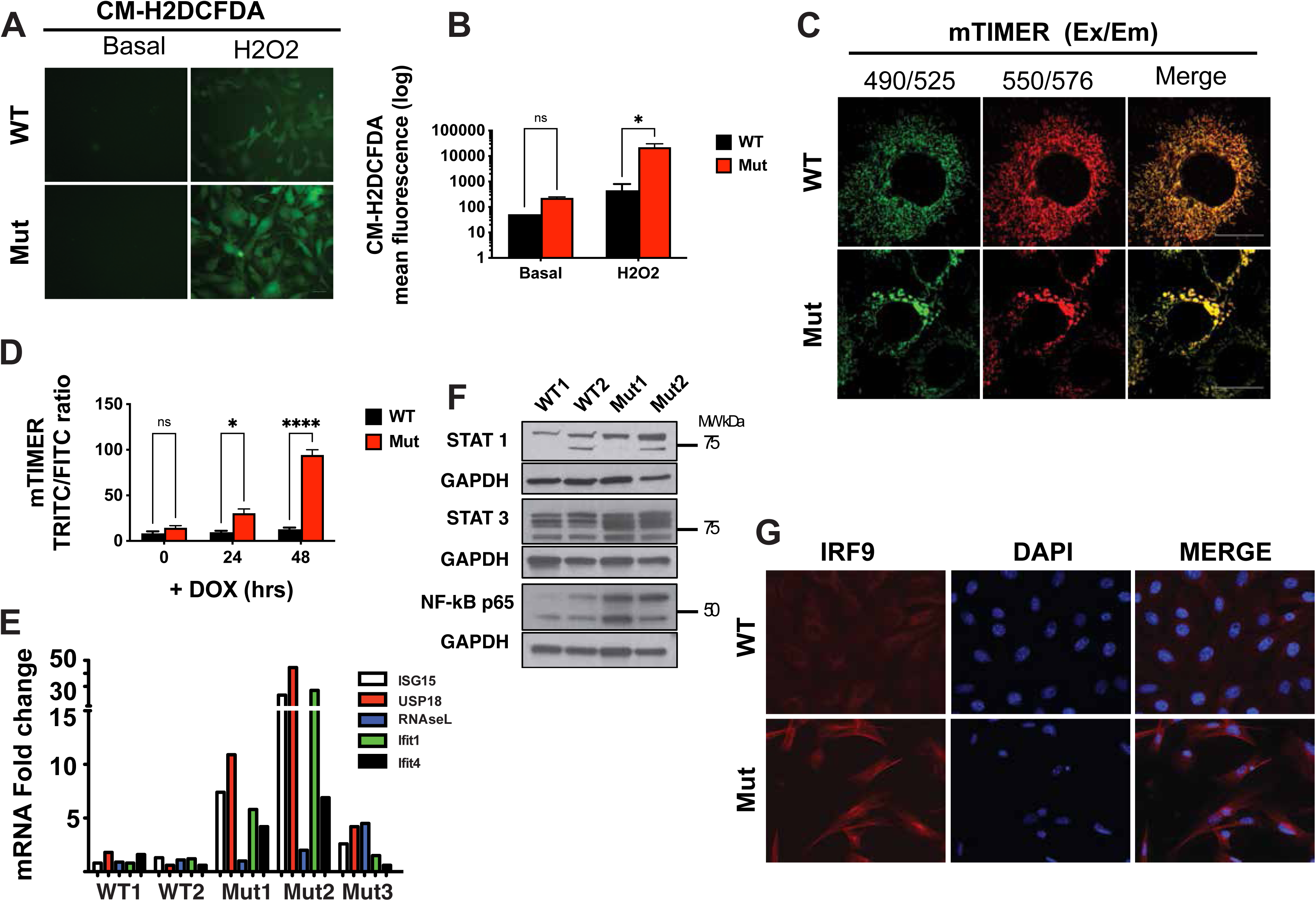
Mitochondrial dysfunction induces oxidative stress and triggers innate immunity in primary-derived fibroblasts. **(A)** Representative images of Mut and WT-derived fibroblasts incubated with the oxidative stress indicator CM-H2DCFDA under basal conditions (left) and treated with 100 μM hydrogen peroxide (H_2_0_2_) for 24 h (right). **(B)** CM-H2DCFDA quantification by flow cytometry (FACS) in Mut (red) and WT (black) fibroblasts under basal conditions and 24 h post-treatment with 100 μM hydrogen peroxide (H_2_0_2_). Bars represent the mean of two different experiments performed in triplicate. Significances were determined by two-way ANOVA and Sidak’s multiple comparison test with \**p*<0.05. **(C)** mTIMER fluorescence in Mut and WT fibroblasts at 490/525 nm (excitation/emission, green, left panel) and 550/576 nm (excitation/emission, red, middle panel). **(D)** FACS analysis of green to red conversion of mTIMER in Mut as compared to WT fibroblasts. mTIMER expression was induced by doxycycline (+DOX) and assessed at 0, 24, and 48 h. Significances were determined by two-way ANOVA and Sidak’s post-comparison test with * p< 0.05 and ****p < 0.0001. **(E)** qPCR measuring the fold-change of five inflammatory genes in three mtDNA mutator (Mut) and two WT-derived fibroblast cell lines: ISG15 (white), USP18 (red), RNAseL (blue), Ifit1 (green). and Ifit4 (black). **(F)** Representative Western blot to assess the level of STAT 1, STAT 3, and NF-KB p65 in Mut and WT-derived fibroblasts (two different cell lines per group). GAPDH was used as loading control. **(G)** Immunofluorescence showing IRF9 expression and localization in Mut and WT-derived fibroblast cell lines. Nuclei are stained with DAPI.

### Mitochondrial Dysfunction and Inflammation in mtDNA Mutator-Derived Fibroblasts

To determine whether mtDNA mutator-derived fibroblasts could serve as a suitable model for dissecting the underlying molecular mechanisms of mitochondrial dysfunction-induced inflammation, we initiated our investigation by assessing the expression of 5 interferon-induced genes (ISG15, USP18, RNAseL, Ifit1, and Ifit4) in mtDNA mutator cells, as compared to WT cells using real-time PCR. Our data revealed an overall higher expression of these genes in mtDNA mutator-derived fibroblasts compared to WT, with 2 out of 3 cell lines (Mut1 and 2) exhibiting a robust induction of these genes (Fig. 8E). Furthermore, mtDNA mutator-derived cells demonstrated elevated levels of proteins associated with an immune response, including STAT-1, -3 and NFκB-p65, as confirmed by Western blot analysis (Fig. 8F), and IRF-9, as determined by immunofluorescence (Fig. 8G). Collectively, these findings indicate an up-regulation of inflammatory genes and proteins in mtDNA mutator-derived cells, establishing them as a suitable model for studying the molecular pathways activated by mitochondrial dysfunction to induce inflammation.

**Figure 8.**
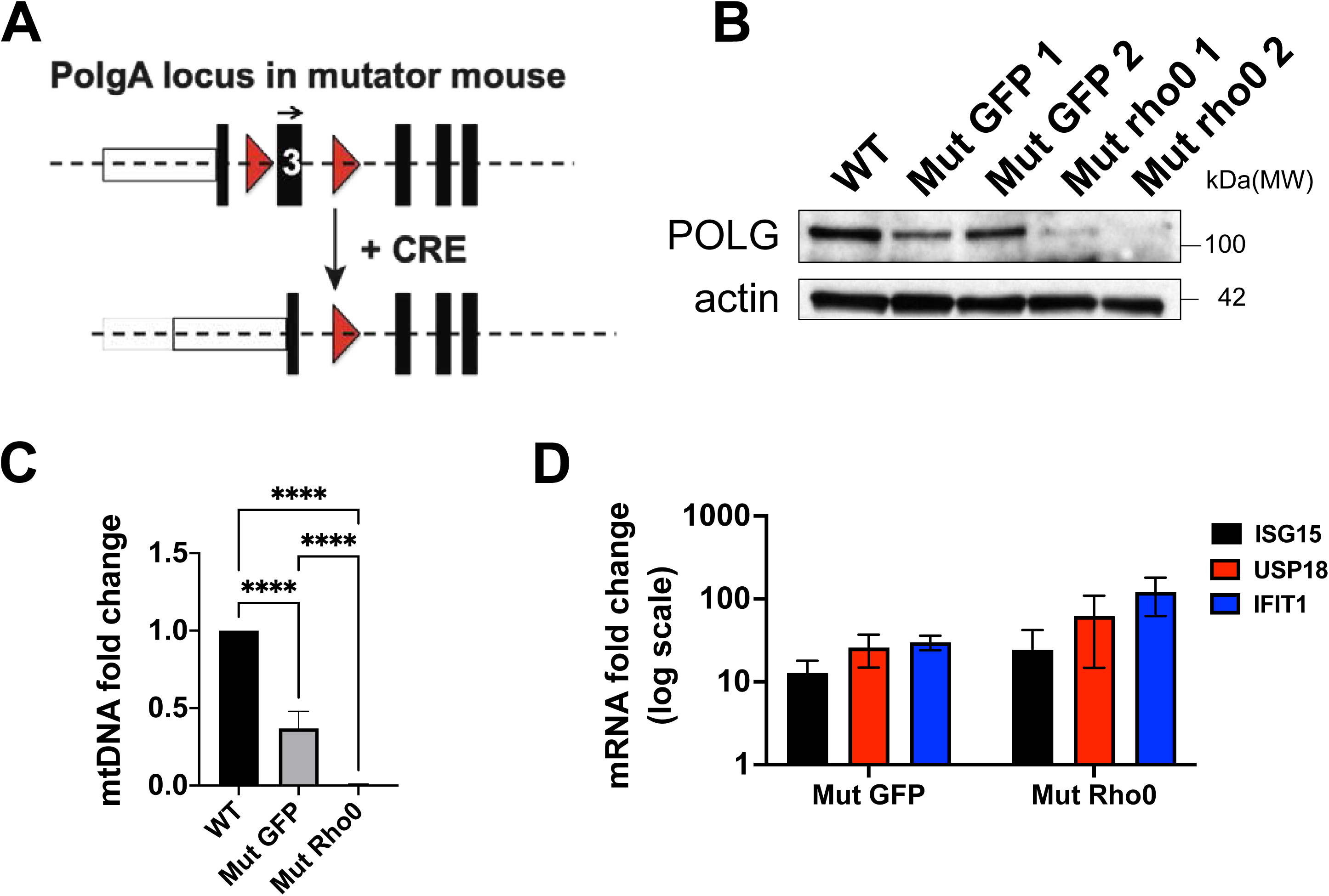
mtDNA is not required to trigger the innate response in primary cells. **(A)** Schematic of the *PolgA* locus in the mtDNA mutator mouse. Two loxP sequences (red triangles) are located on both sides of exon 3. Expression of CRE recombinase induces excision of exon 3 leading to *PolgA* knock-out. **(B)** Representative Western blot showing POLGA protein levels in one WT-derived fibroblast cell line, two Mut cell lines expressing GFP (Mut GFP), and two Mut cell lines expressing CRE recombinase (Mut Rho0). **(C)** mtDNA levels in WT, Mut GFP, and Mut Rho0 cells were assessed by qPCR. Significances were determined with one-way ANOVA and Tukey’s multiple comparison test, ****p < 0.0001. **(D)** mtDNA fold-change of three genes involved in the innate immune response, Isg15 (black), Usp18 (red), and Ifit1 (blue), in Mut GFP and Mut Rho0 cell lines as compared to WT fibroblasts. Bars represent the mean of two cell lines, each tested twice.

Given that the leakage of mtDNA molecules into the cytoplasm is reported to strongly activate interferon responses, we investigated whether this could be the cause for the observed activation of the interferon response in the mtDNA mutator mouse. To test this, we depleted mtDNA mutator-derived cells of their mtDNA content and assessed the effect on the expression of interferon-induced genes. Leveraging the fact that the PolgA gene in the mtDNA mutator mouse is flanked by two LoxP sites (Fig. 8A), we were able to knock-out the PolgA gene to produce Rho0 mtDNA mutator cells by Cre recombination. To this end, cells were transduced with a lentivirus expressing either GFP or Cre recombinase, and the level of PolgA protein was assessed by Western blot. As shown in Figure 8B, the level of POLGA protein was almost undetectable in mtDNA mutator cells transduced with CRE after two weeks post-infection, compared to those transduced with GFP or in WT-derived cells. Analysis of mtDNA copy number by real-time PCR confirmed that mtDNA mutator cells transduced with Cre recombinase-expressing virus were significantly depleted of mtDNA content (Fig. 8C). Despite mtDNA depletion, interferon-induced genes (ISG15, USP18, and Ifit1) upregulation in mtDNA mutator fibroblasts remained induced, suggesting that mitochondrial dysfunction induction of interferon response is independent of mtDNA content in this model (Fig. 8D).

In summary, our data demonstrate that mitochondrial dysfunction in mice, due to the accumulation of mtDNA mutations, induces liver steatosis and inflammation, making the mtDNA mutator mouse model suitable for studying the role of mitochondrial dysfunction in NAFLD onset and progression.

## Discussion

This study establishes a causal link between mitochondrial dysfunction and NAFLD development. The phenotypic changes in mtDNA mutator mice, including early-onset lipid droplet accumulation in liver that worsens as mitochondrial dysfunction progresses, reflect the complex role of mitochondria in NAFLD pathophysiology (7,11). The liver, highly dependent on mitochondrial integrity, is particularly vulnerable to dysfunction caused by mtDNA mutations. Our findings of liver steatosis and inflammation in Mut mice support growing evidence linking mitochondrial dysfunction to NAFLD, highlighting the relevance of this model in advancing current understanding of the disease (7).

The down-regulation of cytochrome *c* oxidase subunit 1 (MTCO1) and concomitant up-regulation of succinate dehydrogenase B (SDHB) in Mut liver tissue (Fig. 1C and D) reflect distinct alterations within the OXPHOS system. The reduction in Complex IV activity, corroborated by COX histochemical staining (Fig. 1E), underscores the functional consequences of mtDNA mutations on the electron transport chain. Importantly, the selective accumulation of lipid droplets in hepatic tissue highlights a tissue-specific dysregulation of lipid metabolism, reinforcing the mechanistic link between mitochondrial dysfunction and NAFLD pathogenesis (Fig. 1F) (44).

The observed hepatomegaly in Mut mice prompted us to investigate serum lipid metabolism. Consistent with previous studies, triglyceride levels are lower in the mtDNA mutator mouse across all ages (21). Interestedly, our findings show an age-dependent fluctuation in serum cholesterol and phytosterol levels suggesting a complex influence of mitochondrial dysfunction on systemic lipid homeostasis (Fig. 2). Elevated levels of 7α-hydroxy-4-cholesten-3-one (C4) further point to impaired cholesterol clearance through bile acid synthesis (Fig. 2D). Collectively, these findings demonstrate an intricate interplay between mitochondrial dysfunction and cholesterol metabolism (45,46).

The unexpected upregulation of bile acid synthesis-related proteins, such as CYP7A1, together with increased levels of both precursor and cleaved forms of sterol regulatory element-binding protein 1 (SREBP-1), underscores the intricate role of mitochondria in lipid metabolism (Fig. 2A and C). Paradoxically, this SREBP-1 activation coincided with downregulation of key cholesterol synthesis genes, including HMGCR and LDLR, highlighting the complexity of transcriptional and post-transcriptional regulation under mitochondrial dysfunction (Fig. 2F and H). These findings point to a disrupted coordination of lipid metabolic pathways in the setting of NAFLD (47). Activation of SREBP-1 under fat-rich conditions suggests the involvement of regulatory mechanisms beyond the classical cholesterol-mediated pathway (48). Our proteomic analysis revealed that two major negative regulators of SREBP-1, INSIG-1 and INSIG-2, are down-regulated in liver from mtDNA mutator mice, supporting the observed up-regulation of SREBP-1. One possible explanation is that mitochondrial dysfunction induces ER stress, which in turn activates SREBP-1 through various mechanisms, including inhibition of INSIG protein translation (49). Interestingly, genes involved in the cholesterol biosynthetic pathway are concurrently down-regulated in mtDNA mutator mice, a paradox that suggests SREBP-1 activation in this context may be driven more by the inflammatory response than by lipid metabolic needs. In support of this, recent studies have identified SREBP-1 as a downstream target of proteins involved in the interferon response, such as STING, which is notably upregulated in our proteomic analysis (50).

Our quantitative proteomic analysis provided a holistic view of the impact of mitochondrial dysfunction on liver proteins, revealing associations with inflammatory processes, mitochondrial dysfunction-related conditions, and NAFLD (Fig. 4). The upregulated proteins identified in our analysis suggest that the liver is mounting a multifaceted response to metabolic stress, inflammation, immune activation, and oxidative damage. This proteomic profile reflects the liver’s adaptive and pathological responses to mitochondrial dysfunction, further implicating these processes in the progression of NAFLD. Up-regulation of proteins involved in lipid metabolism, glucose regulation, and mitochondrial function (e.g., Hk2, Cyp4f18, Pkm2, Acadm) indicate that the liver might be adjusting its metabolic profile. The presence of proteins associated with inflammation (e.g., Tnfaip8l2, Mmp9, Ctsg, Serpinb6a) suggests an inflammatory response occurring in the liver. Proteins such as Hmox1, Ncf2, Sirt3, and Cyp2b9 suggest that the tissue is responding to oxidative stress or increasing detoxification activities. The up-regulation of proteins such as Akr1b7, Sirt3, Ptpn1, and Ctsg implicate the activation of cellular pathways to repair or mitigate damage, possibly from DNA mutations, oxidative stress, or inflammation. The presence of Vcam1, Icam1, and Csf1r suggests that there may be activation of immune cell recruitment, possibly indicating endothelial activation or a response to tissue damage or infection. The presence of proteins like Mmp9, Mmp2, and Plg-RKT indicates that tissue remodeling and extracellular matrix (ECM) breakdown may be occurring, possibly due to fibrosis or chronic liver damage. When we shift our attention to the down-regulated proteins, the data clearly indicate ongoing mitochondrial dysfunction. Indeed, many of those proteins are related to mitochondrial function (e.g., Ndufs3, Ndufa2, Cox5a, Cox6c, Ndufb4) and are involved in OXPHOS and ETC activity. Some down-regulated genes such as Scd1 (Stearoyl-CoA desaturase 1), Dgat2 (Diacylglycerol O-acyltransferase 2), and Agpat2 (1-acylglycerol-3-phosphate O-acyltransferase 2) are involved in lipid metabolism, including fatty acid synthesis and triglyceride formation. Their down-regulation suggests a decrease in lipid biosynthesis, indicating a tendency to decrease lipid content in the tissue. Interestedly, several cytochrome P450 (CYP) enzymes such as Cyp2c29, Cyp2c39, and Cyp2j5 are also downregulated, which could indicate a reduction in the liver’s ability to metabolize and detoxify endogenous and exogenous compounds.

The study of mtDNA mutator-derived fibroblasts can help to explain the cellular changes caused by mtDNA mutations and mitochondrial dysfunction and might help to understand what is happening in the hepatocytes. These cells rely heavily on glycolysis for ATP production and have a disrupted mitochondrial network, showing how mitochondrial dysfunction affects both energy production and cell structure. Interestingly, the continued increase in inflammatory and interferon-related genes, despite the loss of mtDNA, suggests that mitochondrial dysfunction itself, rather than the release of mtDNA into the cytoplasm, is key to starting and keeping inflammation in this model (51,52). In this context, several signaling pathways activated by dysfunctional mitochondria have been shown to trigger inflammation. These include the release of formylated peptides, cardiolipin, and ROS; each of which may contribute to the inflammatory phenotype observed in the mtDNA mutator mouse (53–55).

In conclusion, our study provides valuable insights into the complex relationship between mitochondrial dysfunction and the development of liver steatosis and inflammation, while also situating these findings within the broader context of the current understanding of NAFLD. For future studies, the new inducible mtDNA mutator mouse model we have generated will provide the opportunity to direct mitochondrial mutations specifically to the liver, allowing us to study their consequences on this organ without the confounding effects of other organs or systems being equally affected by mitochondrial dysfunction. This will serve as a robust tool for exploring the molecular mechanisms that link mitochondrial dysfunction to NAFLD pathogenesis, potentially paving the way for targeted therapeutic strategies (56).

## Author Contributions

G.C. and J.M.R. conceived the project and designed experiments; G.C., K.M.K., K.B., M.S. R.B., M.O., I.P-C. and J.M.R. performed the experiments; J.L., I.B., D.A.S., and L.O. contributed analytic tools; J.M.R, G.C., and R.M.B. analyzed data; J.M.R., and G.C. wrote the paper.

## Institutional Review Board Statement

The animal study protocol was approved by the Institutional Review Board of Karolinska Institutet, protocol number N118/10, of Harvard Medical School, protocol number IS00000927, and of the University of Rhode Island, protocol numbers AN1920-020 and AN1920-014.

## Acknowledgments

We thank Margareta Widing, Katrin Wellfelt, Eva Lindqvist, Karin Lundströmer, Karin Pernold, and Anna Lindberg for technical support.

